# Shear stress-induced restoration of pulmonary endothelial barrier function following ischaemia reperfusion injury requires VEGFR2 signalling

**DOI:** 10.1101/2024.01.10.575020

**Authors:** Don Walsh, Daria S. Kostyunina, John Boylan, Paul McLoughlin

## Abstract

Normal physiological shear stress produced by blood flow is sensed by the vascular endothelium and required for the maintenance of both the normal structure and barrier function of the endothelium. Many common, critical illnesses are characterised by periods of abnormally reduced or absent shear stress e.g. haemorrhagic shock, myocardial infarction and pulmonary embolism and are complicated by oedema formation following restoration of normal perfusion (IRI).

We tested the hypothesis that, in lungs injured by a period of ischaemia and reperfusion (IRI), reduced shear stress contributes to increased endothelial barrier permeability and oedema formation following the restoration of perfusion. Furthermore, we examined the role of VEGFR2 as a mechanosensor in the response of the pulmonary endothelium to altered shear stress in this condition.

Following IRI, we perfused isolated ventilated mouse lungs with a low viscosity solution (LVS) or a higher, physiological viscosity solution (PVS) at constant flow to produce differing shear stresses on the endothelium of the intact pulmonary circulation. Lungs perfused with LVS developed pulmonary oedema due to increased endothelial permeability whereas those perfused with PVS were protected from oedema formation by reduced endothelial permeability. This effect of PVS required normal VEGFR2 tyrosine kinase activity but was unaffected by blocking VEGFA binding to the receptor.

These data show for the first time that shear stress has an important role in restoring endothelial barrier function in the pulmonary circulation following injury and have important implications for the treatment of pulmonary oedema in critically ill patients following ischaemia reperfusion injury.

## Introduction

The endothelium of the microvasculature forms the normal semi-permeable barrier that restricts fluid filtration into the surrounding tissues maintaining it at a low rate that is balanced by clearance of interstitial fluid via the lymphatic drainage (Levick and Michel 2010). There are two key components to the barrier, the endothelial glycocalyx and the monolayer of endothelial cells that are joined by junctional complexes between neighbouring cells, both of which are dynamically regulated by stimuli arising in the local microenvironment (Mehta and Malik 2006, Aman, Weijers et al. 2016). Together, the glycocalyx and the endothelium maintain the oncotic gradient across the glycocalyx that is required to prevent excessive water leak into the interstitium (Pries, Secomb and Gaehtgens 2000, van Haaren, VanBavel et al. 2003, Reitsma, Slaaf et al. 2007, Levick and Michel 2010).

Endothelial cells under physiological conditions are subjected to continuous fluid shear stress i.e. the tangential or “dragging” force exerted on the endothelium wall by blood as it flows through a vessel. This shear stress is sensed by the endothelium and regulates important functions of the endothelium such as the production of the vasoactive mediators that are responsible for flow mediated dilatation and the regulation of vascular resistance (Davies 1995, Snow, Markos et al. 2001, Kwon, Noh et al. 2023). Normal physiological shear stress is also required to amintain both the normal structure and barrier function of the endothelium (Seebach, Dieterich et al. 2000, Dekker, Van Soest et al. 2002, Siddharthan, Kim et al. 2007, Walsh, Murphy et al. 2011, Torres Filho, Torres et al. 2013, Zeng and Tarbell 2014, Rowan, Rochfort et al. 2018, Wang, Kostidis et al. 2020, Kostyunina, Rowan et al. 2023).

Many common, critical illnesses are characterised by abnormally reduced shear stress or periods of absent shear stress e.g. low cardiac output conditions such as haemorrhagic shock, haemodilution following resuscitation, myocardial infarction and critical limb ischaemia (Grace 1994, Mongardon, Dumas et al. 2011). These conditions are complicated by impaired endothelial function and oedema formation, which develops following restoration of perfusion by therapeutic interventions, and form an important part of the syndrome of ischaemia reperfusion injury (IRI) (Pak, Sydykov et al. 2017, Kloner, King and Harrington 2018, Hausenloy, Chilian et al. 2019). IRI is also one of the principal determinants of organ failure immediately after organ transplantation (Erasmus, van Raemdonck et al. 2016, Rampes and Ma 2019). Pulmonary oedema resulting from increased endothelial permeability, so called non-cardiogenic pulmonary oedema, frequently complicates conditions characterised by an episode of low or absent blood flow in the lung such as shock, pulmonary embolism and lung transplantation. This is a particularly serious condition because the resultant impairment of gas exchange leads to systemic hypoxaemia. (Collins, Blank et al. 2013, Matthay, Zemans et al. 2019, Borek, Birnhuber et al. 2023). However, the role of reduced shear stress in oedema formation in the lungs following IRI and whether increasing shear stress towards normal values can contribute to restoration of endothelial barrier function in this condition has not previously been examined.

Given that background, we undertook a series of experiments to test the hypothesis that, in lungs injured by IRI, reduced shear stress conditions contributed to increased endothelial barrier permeability and oedema formation following the restoration of perfusion. Furthermore, given the recent demonstration of a key role of PlexinD1-VEGFR2 complex in mechanotransduction in the endothelium, we examined the role of VEGFR2 in the response of the pulmonary endothelium to altered shear stress following IRI (Mehta, Pang et al. 2020).

## Methods

All procedures involving mice were approved by the University College Dublin Animal Ethics Committee (AREC-18-05-McLoughlin) and undertaken following authorisation by the Health Products Regulatory Authority (AE18982.I401). Adult male and female C57 Bl6 mice (10–12 weeks old) were supplied by UCD Biomedical Facility and housed under specific pathogen free conditions with ad libitum access to food and water. The isolated, ventilated, perfused mouse lung was used to assess vascular function, as previously described (Cahill, Rowan et al. 2012, Rowan, Rochfort et al. 2018). Lungs were isolated post-mortem following induction of deep anaesthesia (pentobarbitone sodium, 120 mg/kg I.P.), administration of heparin (1000i.u./kg) and exsanguination. The isolated lungs were placed in a water jacketed chamber maintained at 37°C, ventilated (Rodent Midi-Vent Model 849 Ventilator, product no: 73-4119 (Hugo-Sachs Elektronic-Harvard Apparatus, March, Germany) at constant tidal volume (250 μl, 90 inflations per minute) with 5% CO_2_ balance air and at a constant positive end expiratory pressure (PEEP) of 2.1 cmH_2_O (1.6mmHg). To prevent the development of progressive atelectasis, a recruitment manoeuvre was undertaken every five minutes throughout the experimental protocol by increasing PEEP to 15.0 cmH_2_O (11.5 mmHg) for three inflations. An initial recruitment manoeuvre was performed immediately after tracheal cannulation and then at five-minute intervals throughout the remainder of the protocol. Following cannulation of the pulmonary artery and the left atrium, the lungs were perfused (0.5 - 3.0 ml/minute) while left atrial pressure (LAP) was maintained at 2.0 mmHg in all conditions. Pulmonary arterial pressure (PAP), left atrial pressure (LAP) and airway pressure signals were continuously measured (P75 Blood Pressure Transducer, product no: 73-0020 and Differential Low-Pressure Transducer, product no: 73-3882 respectively, Hugo-Sachs Elektronic-Harvard Apparatus, March, Germany), digitized (220 Hz) and stored for later analysis (AcqKnowledge Data Acquisition 3.8.2 Analysis Software, Biopac Systems Inc, USA).

Lungs were perfused with one of two different solutions. The first was a low viscosity solution (LVS) with a viscosity relative to water of 1.5 (RV 1.5) i.e. a viscosity close to that of normal plasma (Vaya, Simo et al. 2007, Pedersen, Nielsen et al. 2014). The second solution, called the physiological viscosity solution (PVS), had a higher relative viscosity of 2.5. The choice of RV 2.5 as the optimum physiological viscosity solution (PVS) was based on our previous work (Rowan, Rochfort et al. 2018). This viscosity lies within the range of apparent viscosities displayed by normal blood (Fåhræus and Lindqvist 1931, Chien, Usami et al. 1966). LVS consisted of Dulbeccos’s Modified Eagle’s Medium (DMEM, Sigma, Dublin, Ireland, catalog no. D6046) with Ficoll 70kDa (Sigma, Dublin, catalog no. F2878) added (40 g/l) to provide normal oncotic pressure. PVS was composed of DMEM (Sigma, Dublin, Ireland, catalog no. D6046) with Ficoll 70 kDa (Sigma, Dublin, catalog no. F2878) (33.8 g/l) and Ficoll 400kDa (Sigma, Dublin, catalog no. F4375) added (32.5g/l) to increase viscosity. The sum of the molar concentrations of Ficoll 70kDa and Ficoll 400kDa in PVS was equal to the molar concentration of Ficoll 70kDa in LVS. The viscosity of each perfusion solution was measured at 37°C using an Ostwald capillary viscometer.

After insertion of the pulmonary artery cannula and left atrial cannula, perfusion of the lung was initiated at a flow of 0.5ml/min for one minute. Thereafter, the flow was successively increased to 1.0 ml/min, 2.0 ml/min, and 3.0 ml/min at intervals of one minute. Once a flow of 3.0 ml/min was established the lung was perfused for 10 minutes while the venous effluent was discarded; this allowed the clearance of any residual blood from the pulmonary circulation. Experimental lungs were subjected to ischaemia reperfusion injury (IRI) by stopping perfusion following the initial 10 minute period of perfusion. The lungs were exposed to this warm ischaemia for a period of 20 minutes during which the intravascular pressure was maintained at 2mmHg. Following this, flow (3.0 ml/min) was restarted and the circuit was closed i.e. the venous effluent was recirculated for the remainder of the protocol. Perfusion was continued until the lungs became oedematous (Pinsp > 7.5 mmHg), or until 180 minutes had elapsed. Uninjured control lungs were perfused without interruption at a flow of 3ml/min while the venous effluent was discarded. After 30 minutes recirculation of perfusate was commenced i.e. at a time corresponding to the end of the warm ischaemia period in injured experimental lung groups. Perfusion was continued until the lungs became oedematous (Pinsp > 7.5mmHg) or until 180 minutes had elapsed.

Capillary pressure (Pcap) was determined using the double-occlusion technique at end expiration as previously described (Townsley, Korthuis et al. 1986, Cadogan, Hopkins et al. 1999, Rowan, Rochfort et al. 2018). Vascular permeability was assessed by measurement of extravascular leakage of Evan Blue labelled albumin, as previously described (Rowan, Rochfort et al. 2018). At the end of the period of warm ischaemia when recirculation commenced, the perfusion fluid was switched to an identical fluid (LVS or PVS) containing Evans Blue-labelled albumin (0.5g/100 ml, Sigma). At the termination of the experiment, the vasculature was flushed for five minutes with a perfusate that was identical except that it did not contain Evans Blue labelled albumin (3 ml/min) until the draining perfusate was clear, so that only extravasated Evans Blue-labelled albumin remained in the lungs. Wet to dry weight ratios were measured to assess lung fluid content. Formamide (>99.5%, Sigma) was then added to each dried lung and incubated at 70°C for 1 h to extract Evans Blue dye. The lung was then homogenized, the homogenate cleared by centrifugation, and the concentration of Evans Blue determined by absorbance at 620 nm. Tissue content was expressed as micrograms of Evans Blue per milligram of lung dry weight (Yin, Michalick et al. 2016, Rowan, Rochfort et al. 2018). The change in Heparan sulphate in the perfusate during the period from the end of the warm ischaemia until the end of the protocol was measured by ELISA (LS-Bio Massuchusets, USA, Catalogue No. LS-F39210) according to the manufacturer’s instructions as an index of glycocalyceal breakdown and shedding during that period (Annecke, Fischer et al. 2011, Schmidt, Yang et al. 2012).

At selected points throughout the experimental protocols, mean airway and vascular pressures were determined immediately before a regular recruitment manoeuvre or just prior to the last recruitment manoeuvre before termination of the experiment. Mean peak inspiratory pressure (Pinsp) was calculated as the mean of the peak airway pressure values from 10 consecutive inflations and mean positive end expiratory pressure (PEEP) as the mean value of ten consecutive measurements at end expiration during the same respiratory cycles. In some experimental series, the rate of change of Pinsp in each lung was calculated as Pinsp at the end of the protocol less Pinsp 10 minutes after perfusion recommenced following the warm ischaemia period (when Pinsp was stable) divided by the interval between those two measurements. Mean pulmonary arterial pressure (PAP) was calculated as the mean of 10 values of pulmonary arterial pressure measured at end expiration during the same 10 consecutive respiratory cycles. Mean left atrial pressure was calculated using 10 measurements of LAP taken at the same time points.

Mean pulmonary vascular resistance (PVR) under experimental conditions was calculated using the following conventional formula:

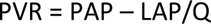

Where PAP is mean pulmonary arterial pressure, LAP is mean left atrial pressure and Q is perfusate flow rate. The resistance to flow in a vascular bed is a function of both the viscosity of the perfusing fluid and geometry of the vascular bed. The contribution of vascular geometry to flow resistance is termed vascular hindrance (Chatpun and Cabrales 2010, Hoffman 2011, Rowan, Rochfort et al. 2018). Since we perfused different groups of lungs with fluids of differing viscosities, we wished to obtain an index of vascular hindrance that was independent of the perfusing fluid viscosity and thus allowed comparison of vascular hindrance between groups. To do this we calculated the resistance that would have been measured had the lungs been perfused with solutions whose viscosities equalled that of water as follows:

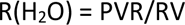

Where RV is the relative viscosity of the perfusate. It is worth noting that the geometric properties that determine hindrance can vary acutely e.g., due to vasoconstriction or vascular recruitment (Chatpun and Cabrales 2010, Rowan, Rochfort et al. 2018).

Statistical analysis was performed using GraphPad Prism software version 9.0 (GraphPad Software, San Diego, CA, USA). Values of n give the number of separate lungs in each group. Data are presented as means (SD). Statistical significance of the difference between means was determined using two-tailed, unpaired *t*-tests. When multiple comparisons of means were undertaken, the Holms-Sidak stepdown correction was used (Ludbrook 1998). The duration of survival of lungs in each group (i.e. time until oedema developed or the end of the protocol was reached) was calculated using the Kaplan-Meier product limit estimate and the significance between these as determined using the log-rank test. When more than two curves were compared in a series of experiments, the Holms-Sidak stepdown correction was used (Ludbrook 1998). In all experiments, a value of P<0.05 was considered statistically significant; where P>0.001, the exact P value is shown.

## Results

### Physiological viscosity solution restored endothelial barrier function following ischaemia reperfusion injury

To determine the effects of different viscosity perfusion solutions on oedema formation, lungs were randomly assigned to three different groups (Figure 1). Figure 1A shows the record of an uninjured control lung in which perfusion with LVS was continued uninterrupted until the end of the protocol (180 minutes) without evidence of oedema formation. Figure 1B shows the record of a lung from the LVS IRI group in which, following an initial period of stabilisation, perfusion was stopped for 20 minutes and then recommenced i.e. an episode of warm ischaemia-reperfusion injury. During the period of ischaemia, Pinsp rose progressively but then recovered to the initial baseline value following reinstitution of perfusion. Subsequently, Pinsp increased progressively until it exceeded the critical value of 7.5mmHg indicating the presence of severe oedema. Figure 1C shows the record from a lung perfused with PVS and then subjected to IRI. During the period of ischaemia, Pinsp increased slowly and progressively in a similar manner to the LVS IRI lung. Following reperfusion Pinsp fell to the initial level seen before the ischaemic period and continued for 180 minutes without reaching the critical Pinsp threshold of 7.5mmHg.

**Figure 1.**
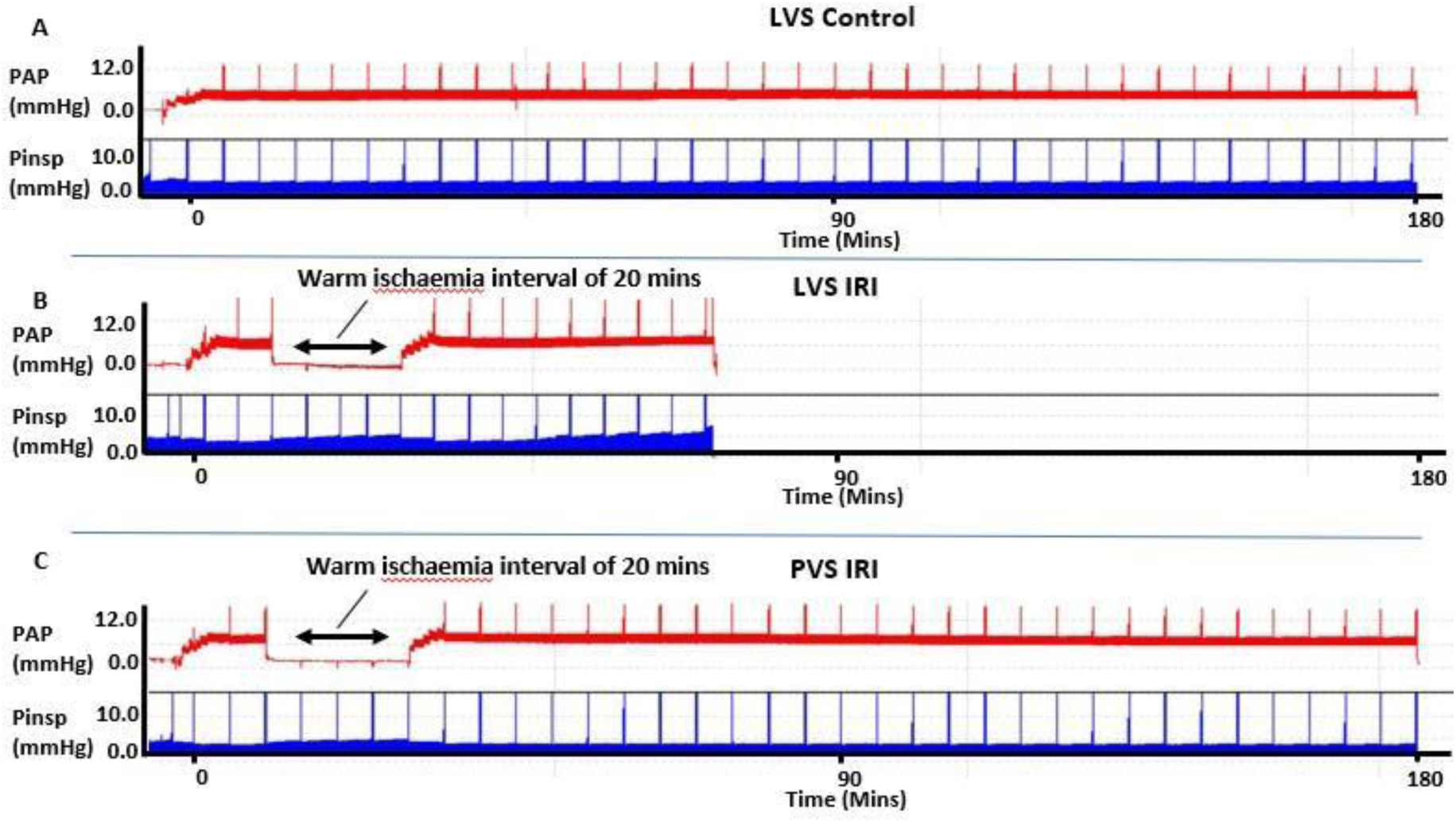
Experimental protocol illustrated using original experimental records. Copies of sample experimental records showing continuous measurements of pulmonary arterial pressure and airway pressure throughout perfusion protocols. A, recording of pulmonary arterial and airway pressures from an uninjured control lung perfused with LVS which completed 180 minutes as Pinsp remained <7.5mHg. B, recording of pulmonary arterial pressure and airway pressures from a lung perfused with LVS in which warm ischaemia injury was produced by stopping flow for 20 minutes. Following recommencement of perfusion (LVS IRI), Pinsp increased progressively to 7.5mmHg indicating critical oedema formation and the experiment was stopped at this point. C, recoding of pulmonary arterial and airway pressures in a lung perfused with physiological viscosity solution. Following cessation of flow to produce warm ischaemia and subsequent reperfusion (PVS IRI), the protocol was continued for 180 minutes as Pinsp remained stable and did not increase to 7.5 mmHg. Note the spike artefact on the pressure records is caused by intermittent regular recruitment manoeuvres of lung to prevent progressive atelectasis. IRI, ischaemia reperfusion injury. LVS, low viscosity solution. Pinsp, peak inspiratory pressure. PAP, pulmonary arterial pressure. PVS, physiological viscosity solution.

In the LVS IRI and PVS IRI groups, mean Pinsp had increased significantly at the end of the period of warm ischaemia injury compared to the mean values observed prior to the ischaemic period demonstrating a reduction in dynamic compliance that was compatible with the onset of early pulmonary oedema (Table 1). This was not seen in the LVS Con group in which Peak inspiratory pressures remained unchanged during the corresponding period (Table 1). Following restoration of perfusion, peak inspiratory pressures initially recovered to baseline values in the two groups subjected to IRI (Table 1).

**Table 1.** Mean (SD) peak airway pressures measured immediately before the period of warm ischaemia, at the end of the period of ischaemia and 10 minutes after restoration of perfusion.

|  | Pre-ischaemia | Ischaemia | Post-ischaemia |
| --- | --- | --- | --- |
| **LVS Con (n=10)** | 4.6 (0.4) | 4.5 (0.3) | 4.5 (0.3) |
| **LVS IRI (n=10)** | 4.6 (0.4) | 5.11 (0.3)* | 4.7 (0.3) |
| **PVS IRI (n=10)** | (4.6 (0.4) | 5.1 (0.2)* | 4.6 (0.2) |

Values presented as mean (SD). LVS Con, lungs perfused with low viscosity solution not subjected to warm ischaemia injury. LVS IRI, lungs perfused with low viscosity solution following warm ischaemia injury. PVS IRI, Lungs perfused with optimum physiological viscosity solution following warm ischaemia injury. Pre-ischaemia, peak airway pressures immediately before warm ischaemia injury. Ischaemia, peak airway pressures after 20 minutes of warm ischaemia immediately prior to recommencing perfusion. Post-ischaemia, peak airway pressures following 10 minutes of reperfusion following warm ischaemia injury. In the LVS con group measurements were made at the corresponding time points although perfusion was continued without interruption. \* indicates significant difference ( $P < 0.001$ ) from the Pre-ischaemia injury value in that group.

Lung survival was significantly shorter in the LVS IRI group than in the control group confirming lung injury as a result of the warm ischaemia period (Figure 2A). In contrast, in the PVS IRI group lung survival times were significantly greater than in the LVS IRI group and not significantly different from the LVS control lungs (Figure 2A). Interestingly, at the end of the protocol, mean PAP in all three groups was similar (Figure 2B) despite the very different viscosities of the perfusates used.

**Figure 2.**
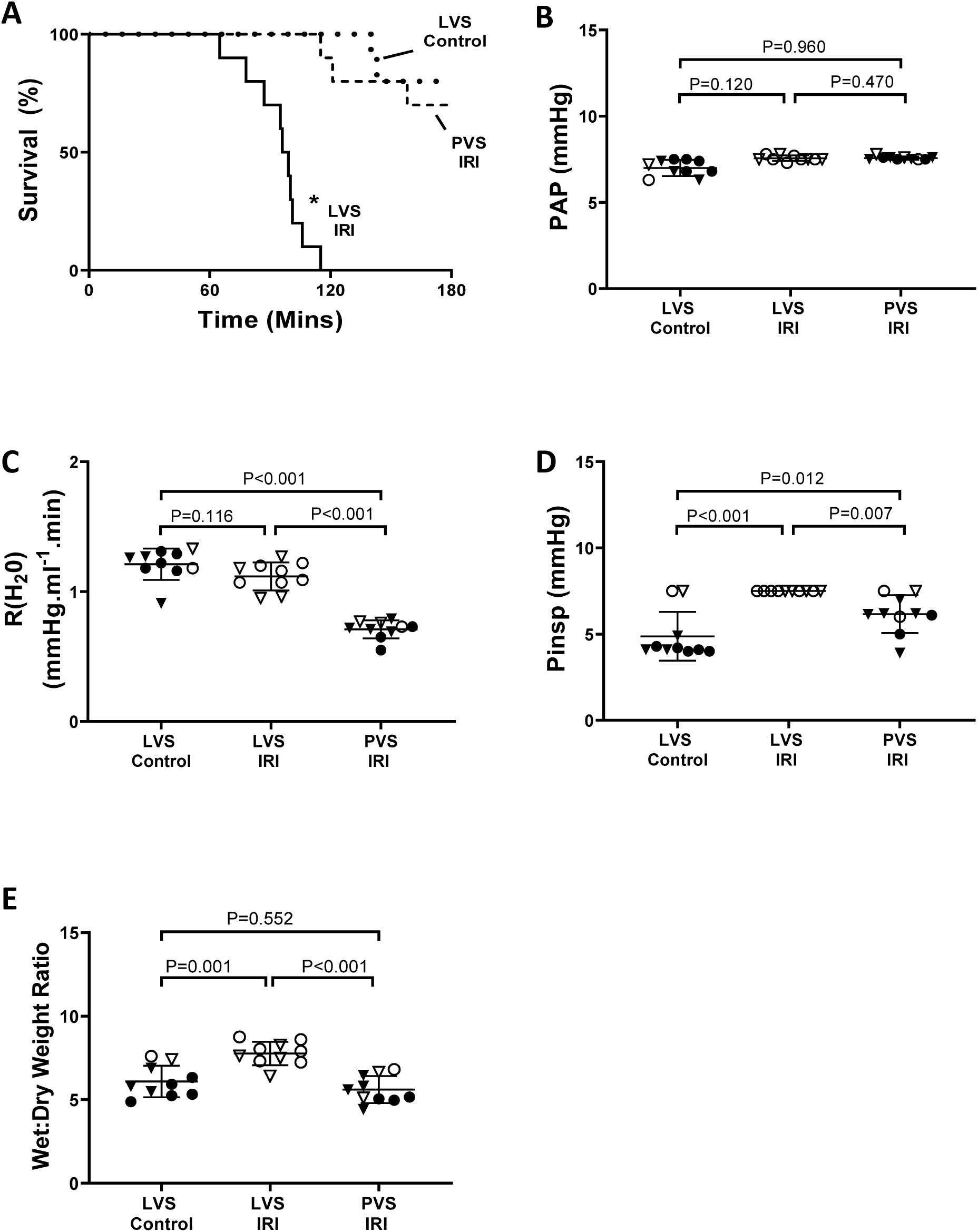
Effect of PVS on lung survival following ischaemia reperfusion injury. A, Kaplan-Meier survival curves for uninjured, control lungs perfused with low viscosity solution (LVS Control, n=10), injured lungs perfused with LVS (LVS IRI, n=10) and injured lungs perfused with physiological viscosity solution (PVS IRI, n=10). * indicates statistically significant difference from the two other groups (P<0.001, Log rank test with Holm-Sidak correction for multiple comparisons). B, mean (SD) pulmonary artery pressure (PAP) in each group at termination of perfusion (n=10 in each group). C, mean (SD) R(H_2_0) in groups at termination of perfusion (n=10 in each group). D, mean (SD) airway pressure (Pinsp) in each group at termination of perfusion (n=10 in each group). E, mean (SD) wet to dry weight ratio in each group at termination of perfusion (n=10 in each group). Open symbols represent lungs in which perfusion was terminated due to the development of oedema (Pinsp>7.5mmHg) before 180 minutes. Filled symbols represent lungs in which oedema did not develop. Triangular symbols represent male lungs and circular symbols represent female lungs. Exact P values were computed using an un-paired t-test followed by Holm-Sidak correction for multiple comparisons. IRI, ischaemia reperfusion injury. LVS, low viscosity solution. Pinsp, peak inspiratory pressure. PAP, pulmonary arterial pressure. PVS, physiological viscosity solution. R(H_2_O), pulmonary vascular resistance corrected to perfusion with a solution whose relative viscosity equals one.

Determination of R(H_2_O) showed that PAP was not elevated in the PVS group as a result of a marked (P<0.001) reduction of R(H_2_O) when compared to both the LVS Control and LVS IRI groups (Figure 2C). At the end of their perfusion periods, Pinsp in the LVS control and PVS IRI were significantly less than in the LVS IRI group (Figure 2D) and, in keeping with this, the wet:dry weight ratios of both the LVS control and PVS IRI groups were significantly less than those of the LVS IRI group (Figure 2E), confirming that improved lung survival in the PVS group was due to a reduction in oedema formation.

### PVS reduces endothelial barrier permeability following IRI

In a further series of experiments the effect of PVS on endothelial barrier permeability following IRI was examined. Lungs were allocated to one of the following experimental protocols in groups of three (triplets): LVS control, LVS IRI or PVS IRI. In each triplet, a first lung was perfused with LVS, subjected to IRI and following reperfusion the protocol was continued until Pinsp reached >7.5 mmHg. Thereafter, in a randomised order, an uninjured lung was perfused with LVS (LVS control) and an injured lung with PVS (PVS IRI) was perfused for a period matching that of the LVS IRI lung or until Pinsp reached >7.5 mmHg, whichever occurred first. Thus, for every lung in the LVS IRI group there was an LVS control lung and a PVS IRI lung perfused for an identical period. In all three groups PAP was similar (Figure 3A), while R(H_2_O) was significantly lower in the PVS IRI group than in the other two groups (Figure 3B). Oedema formation in the PVS IRI group was significantly reduced when compared to the LVS IRI group, as demonstrated by a lower rate of increase in Pinsp (Figure 3C). The extravasation of Evans Blue-labelled albumin was significantly reduced in the PVS IRI group when compared with LVS IRI group (Figure 3E), demonstrating that PVS had a protective effect on endothelial barrier function following IRI. Interestingly, the extravasation of Evans Blue-labelled albumin was similar in the PVS IRI and LVS Control groups suggesting that PVS completely reversed the damaging effect of IRI on barrier function. The change in heparan sulphate concentration in the perfusate from the recommencement of perfusion until the end of the protocol was measured as an index of glycocalyceal injury during that period (Annecke, Fischer et al. 2011, Schmidt, Yang et al. 2012). This was reduced in the PVS IRI group when compared to the LVS IRI group, which is keeping with the reduced capillary permeability observed during perfusion with PVS (Figure 3F).

**Figure 3.**
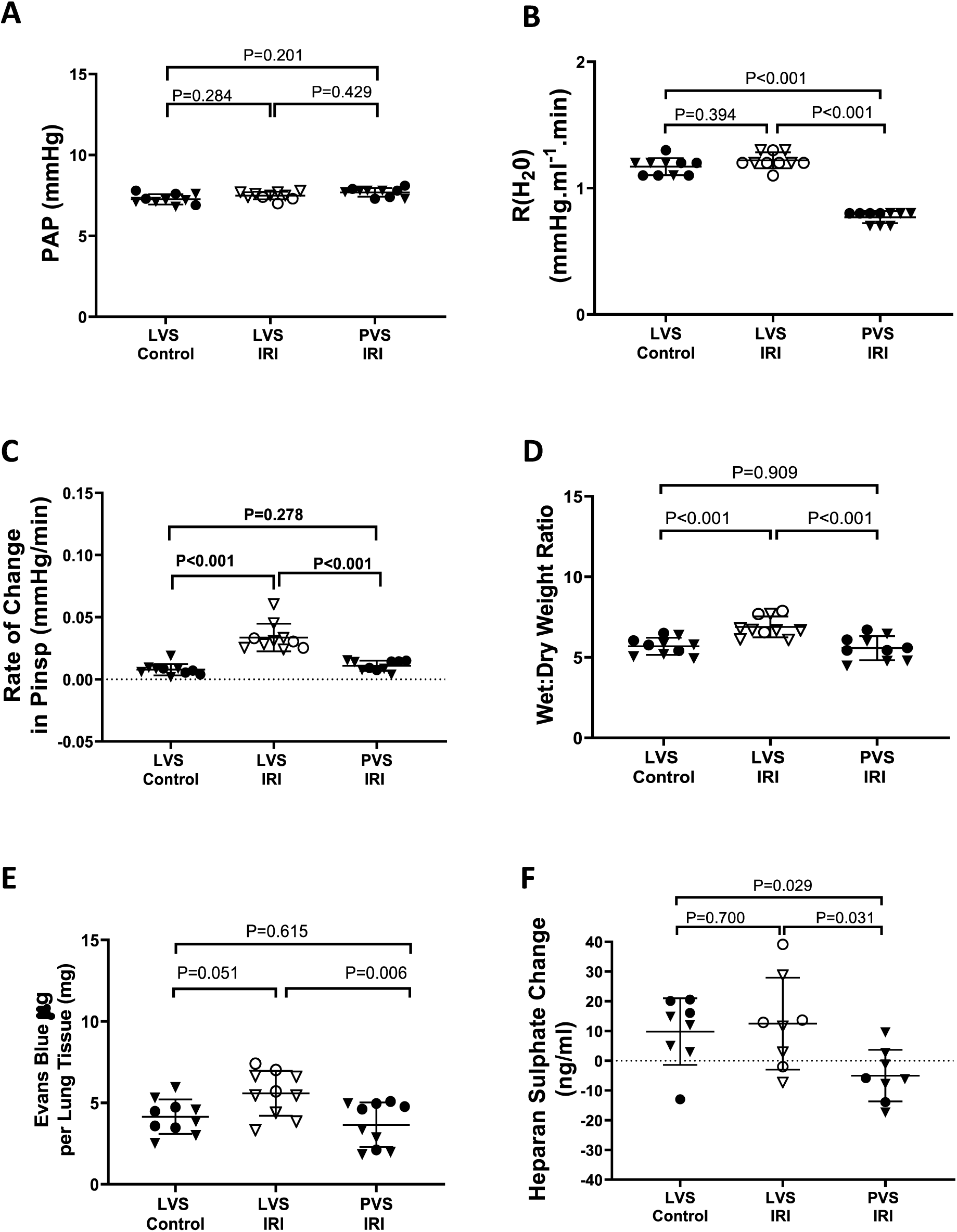
Effect of PVS on endothelial barrier function. A, mean (SD) pulmonary artery pressure (PAP) in the three different groups at termination of perfusion (n=10 in each group). B, mean (SD) R(H_2_0) in groups at termination of perfusion (n=10 in each group). C, mean (SD) rate of change of airway pressure (Pinsp) groups throughout period of perfusion (n=10 in each group). D, mean (SD) wet to dry weight ratio in groups at termination of perfusion (n=10 in each group). E, mean (SD) Evans blue-labelled albumin concentration in lung tissue at termination of perfusion (n=10 in each group). F, Change in heparan sulphate concentrations in the perfusate in each of the three groups following the period of warm ischaemia injury (n=8 in each group). Open symbols represent lungs in which perfusion was terminated due to the development of oedema (Pinsp>7.5mmHg) before 180 minutes. Filled symbols represent lungs in which oedema did not develop. Triangular symbols represent male lungs and circular symbols represent female lungs. Exact P values were computed using an un-paired t-test followed by Holm-Sidak correction for multiple comparisons. IRI, ischaemia reperfusion injury. LVS, low viscosity solution. Pinsp, peak inspiratory pressure. PAP, pulmonary arterial pressure. PVS, physiological viscosity solution. R(H_2_O), pulmonary vascular resistance corrected to perfusion with a solution whose relative viscosity equals one.

### The protective effect of PVS on endothelial barrier function depends on normal VEGFR2 signalling

VEGFR2 is an essential component of a mechanosensory and transduction mechanism in endothelial cells incorporating PLXND1 as a direct force sensor together with the co-receptor neuropilin 1 (Jin, Ueba et al. 2003, Lee and Koh 2003, Coon, Baeyens et al. 2015, Mehta, Pang et al. 2020). Since increasing the viscosity of a Newtonian fluid at constant flow must increase shear stress on the vessel wall, we postulated that the protective effect of PVS on endothelial barrier function was through activation of the mechanosensing function of VEGFR2 and autophosphorylation, the first step in its downstream signalling pathway (Mehta, Pang et al. 2020). Therefore, we examined the effect of SU1498 (10 micromole/L), a potent and selective inhibitor of VEGFR2 autophosphorylation (Strawn, McMahon et al. 1996).

Following random allocation of mice, lungs were perfused in time-matched sets of four. An initial lung was perfused with LVS, injured by warm ischaemia as previously described (Figure 1), following which perfusion was recommenced and continued until Pinsp exceeded 7.5mmHg. Subsequently, one lung was perfused following injury with PVS alone (PVS-IRI), another lung with LVS containing the VEGFR2 inhibitor (LV-IRI SU1498) and another with PVS containing the VEGFR2 inhibitor (PVS-IRI SU1498) for an identical period (Figure 4). In all four groups PAP remained closely similar (Figure 4A). R(H_2_O) in both the PVS IRI and PVS IRI SU1498 groups was significantly lower (P<0.001) than in the LVS IRI and LVS IRI SU1498 groups (Figure 4B), an effect similar to that seen in the previous experiments (Figure 3). Perufusion with PVS reduced significantly the rate of oedema formation compared to LVS as demonstrated by a reduced rate of increase of Pinsp and a reduced wet:dry ratio (Figure 4C and 4D), confirming our previous findings (Figure 3). Furthermore, perfusion with PVS reduced leak of albumin compared to LVS suggesting that its protective effect was produced by improving endothelial barrier function (Figure 4E). Addition of SU1498 abolished the protective effect of PVS in IRI as demonstrated by a significant increase in the rate of increase in Pinsp seen in the PVS IRI SU1498 group compared to the PVS IRI group without added VEGFR2 inhibitor (Figure 4C) and a significantly higher wet:dry weight ratio (Figure 4D). The addition of SU1498 to the perfusate also caused a significant increase in the leak of Evans blue labelled albumin when compared to the PVS IRI group demonstrating that the beneficial effect of PVS on endothelial barrier function required VEGFR2 mediated signalling (Figure 4E). Interestingly, in the presence of VEGFR2 inhibition, the rate of increase of Pinsp, wet:dry ratio and albumin leak were similar in LVS perfused lungs (LVS IRI SU1498) and in the PVS perfused lungs (PVS IRI SU1498) suggesting that signalling through VEGFR2 accounted completely for the protective effects of PVS in the presence of endothelial injury (Figure 4C, 4D and 4E).

**Figure 4.**
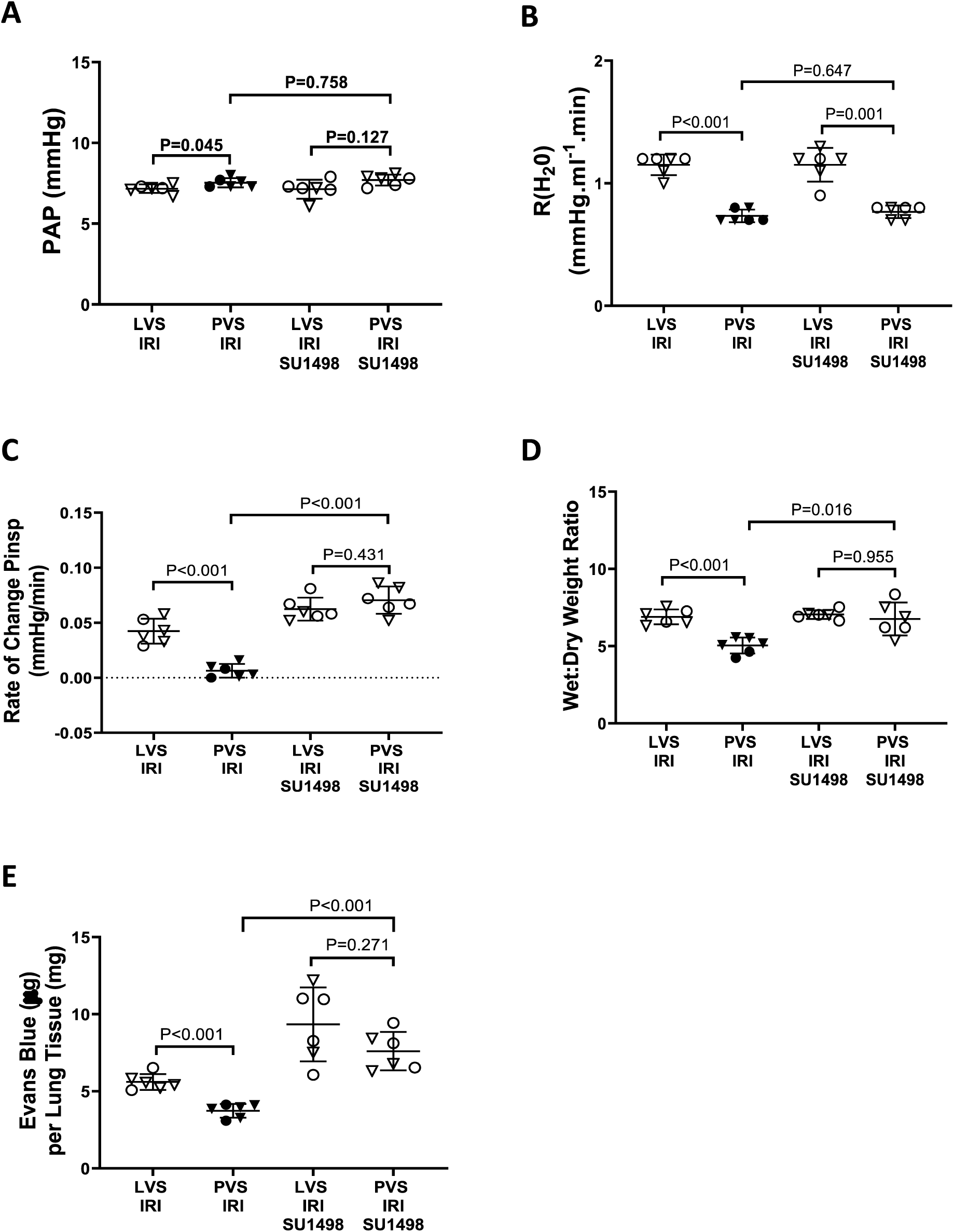
The protective effect of PVS depends on normal VEGFR2 signalling. A, mean (SD) pulmonary artery pressure (PAP) in the four different groups at termination of perfusion (n=6 in each group). B, mean (SD) R(H_2_0) in groups at termination of perfusion (n=6 in each group). C, mean (SD) rate of change of airway pressure (Pinsp) groups throughout period of perfusion (n=6 in each group). D, mean (SD) wet to dry weight ratio in groups at termination of perfusion (n=6 in each group). E, mean (SD) Evans blue-labelled albumin concentration in lung tissue at termination of perfusion (n=6 in each group). Open symbols represent lungs in which perfusion was terminated due to the development of oedema (Pinsp>7.5mmHg) before 180 minutes. Filled symbols represent lungs in which oedema did not develop. Triangular symbols represent male lungs and circular symbols represent female lungs. Exact P values were computed using an un-paired t-test followed by Holm-Sidak correction for multiple comparisons. IRI, ischaemia reperfusion injury. LVS, low viscosity solution. Pinsp, peak inspiratory pressure. PAP, pulmonary arterial pressure. PVS, physiological viscosity solution. R(H_2_O), pulmonary vascular resistance corrected to perfusion with a solution whose relative viscosity equals one. SU1498, VEGFR2 tyrosine kinase inhibitor.

### The protective effect of PVS is independent of change in capillary pressure

Although our results had shown that PVS reduced oedema formation by reducing endothelial barrier permeability, we wished to examine the possibility that it also reduced oedema formation by reducing capillary hydrostatic pressure. To do this we undertook further experiments in which we measured capillary pressure after the lungs had stabilised following the recommencement of perfusion. We used a time matched design identical to that described above in which, following random allocation, lungs were perfused in time-matched sets of four (Figure 4). In this series of experiments, we also modified the injury protocol to examine the possibility that the presence of different perfusates during the period of warm ischaemia could have modified the injury and thus contributed to the different degrees of injury in the different groups. All lungs were perfused with LVS from initiation of the experimental perfusion protocol until the period of warm ischaemia injury was completed. Following this the perfusion solution was switched to the test solution i.e. control solution (LVS), PVS, LVS with SU1498 added or PVS with SU1498 added.

PAP was similar in all four groups at the end of the perfusion protocol (Figure 5A). PVS reduced vascular hindrance in the injured lungs similarly, whether or not SU1498 was present (Figure 5B). PVS reduced the rate of increase of Pinsp following IRI and this protective effect was abolished by blockade of VEGFR2 (Fig. 5C). PVS also reduced the wet:dry weight ratio at the end of the protocol confirming that it attenuated oedema formation (Figure 5D) and its protective effects were abolished by VEGFR2 blockade with SU1498 (Figure 5C and 5D). An improvement in endothelial barrier function was again observed, as demonstrated by significantly reduced leak of Evans blue labelled albumin in the PVS IRI group, and this action was lost when VEGFR2 signalling was blocked (Figure 5E). Importantly, perfusion with PVS did not alter capillary hydrostatic pressure following IRI when compared to LVS IRI and the presence of SU1498 did not alter capillary pressure significantly regardless of the perfusate (Figure 5F). Thus, the attenuation of oedema formation produced by PVS was not due to a change in capillary filtration pressure, suggesting that the protective effect was mediated by its beneficial action on endothelial barrier function.

**Figure 5.**
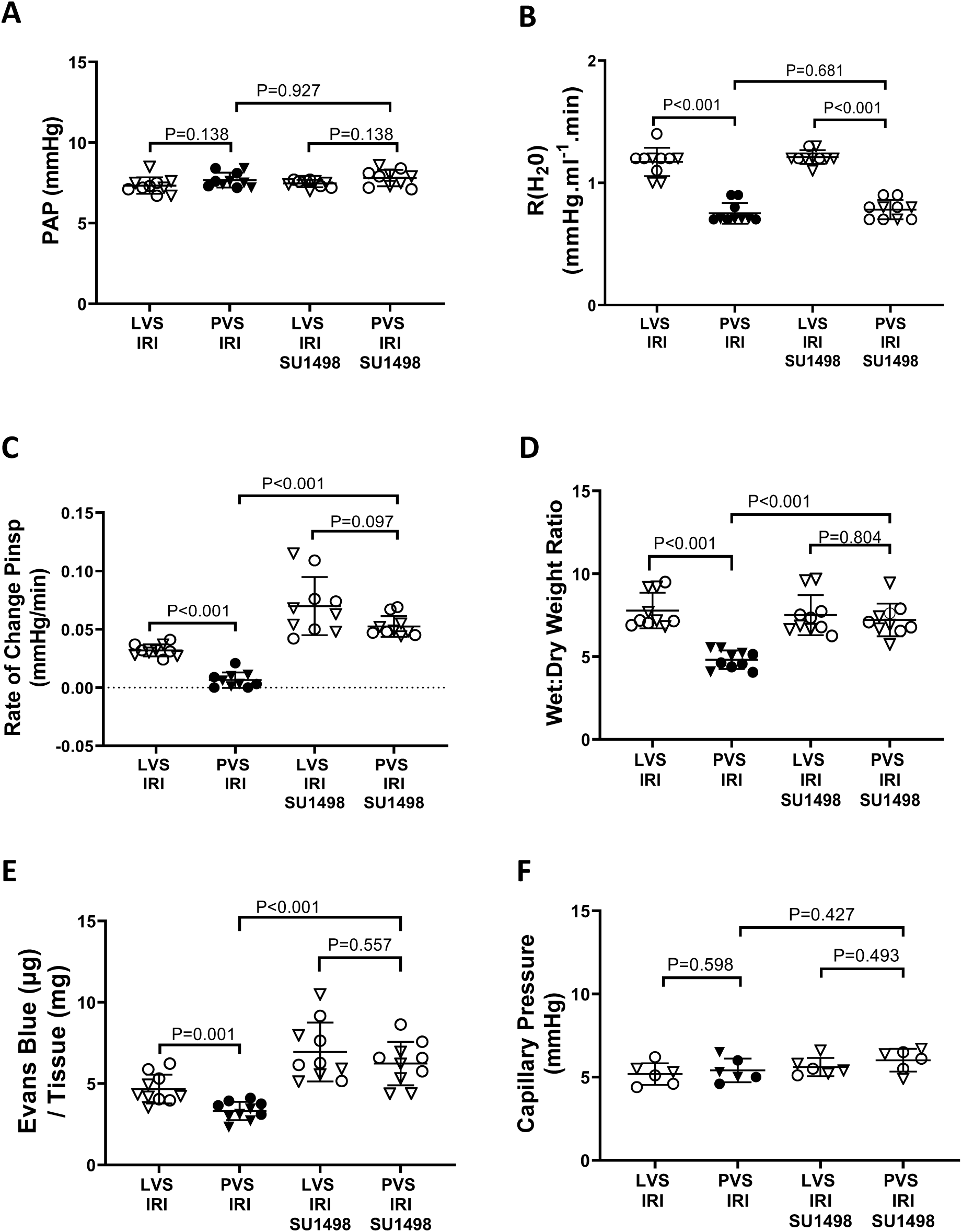
Protective effect of PVS is independent of change in capillary pressure. A, mean (SD) pulmonary artery pressure (PAP) in the four different groups at termination of perfusion (n=10 in each group). B, mean (SD) R(H_2_0) in groups at termination of perfusion (n=10 in each group). C, mean (SD) rate of change of airway pressure (Pinsp) groups throughout period of perfusion (n=10 in each group). D, mean (SD) wet to dry weight ratio in groups at termination of perfusion (n=10 in each group). E, mean (SD) Evans blue-labelled albumin concentration in lung tissue at termination of perfusion (n=10 in each group). F, mean (SD) capillary pressure in each group following reperfusion (n=6 in each group). Open symbols represent lungs in which perfusion was terminated due to the development of oedema (Pinsp>7.5mmHg) before 180 minutes. Filled symbols represent lungs in which oedema did not develop. Triangular symbols represent male lungs and circular symbols represent female lungs. Exact P values were computed using an un-paired t-test followed by Holm-Sidak correction for multiple comparisons. IRI, ischaemia reperfusion injury. LVS, low viscosity solution. Pinsp, peak inspiratory pressure. PAP, pulmonary arterial pressure. PVS, physiological viscosity solution. R(H_2_O), pulmonary vascular resistance corrected to perfusion with a solution whose relative viscosity equals one. SU1498, VEGFR2 tyrosine kinase inhibitor.

It is worth noting that in the presence of the VEGFR2 inhibitor SU1498 in either the LVS or PVS the rate of oedema formation was increased to such an extent that all those lungs had reached the critical Pinsp (7.5mmHg) between 26 minutes and 75 minutes, which in all cases occurred at a time when the matched LVS IRI and PVIS IRI lungs remained stable. Furthermore, in the presence of VEGFR2 inhibition, the rate of oedema formation and albumin leak were similar in LVS perfused lungs (LVS IRI SU1498) and in the PVS perfused lungs (PVS IRI SU1498) suggesting that signalling through VEGFR2 accounted completely for the protective effects of PVS in the presence of endothelial injury (Figure 5C, 5D and 5E).

### Protective effect of PVS is independent of VEGFA binding to the VEGFR2-neuropillin complex

Binding of VEGF-A to VEGFR2 and its co-receptor neurophilin-1 (NRP-1) increases endothelial permeability (Eliceiri, Paul et al. 1999, Carmeliet 2003). Therefore, we wished to determine if the protective effective of perfusion with PVS depended on VEGFA binding to VEGFR2. To address this issue, we used a small molecule inhibitor that competitively blocks the binding of VEGF-A to the NRP-1 co-receptor site on the NRP-1/VEGFR2 complex (EG00229) and prevents the VEGF-A induced increase in endothelial permeability (Jarvis, Allerston et al. 2010, Fantin, Lampropoulou et al. 2017, Powell, Mota et al. 2018). Following a period of warm ischaemia as already described, lungs were perfused until Pinsp > 7.5 mmHg or 100 minutes had elapsed i.e. a period greater than the maximal period for which any lung had survived in the presence of the inhibitor of VEGFR2 intracellular signalling (SU1498).

Lungs were randomly allocated to three groups following initial perfusion with LVS until a 20-minute period of warm ischaemia was completed. Following this, the perfusion solution was switched to one of the following: PVS alone (PVS IRI), PVS containing the inhibitor of VEGFR2 tyrosine kinase activity (PVS IRI SU1498) or PVS containing EG00229 (30 micromole/l) to inhibit VEGF-A binding to the VEGFR2-neuropillin complex (PVS IRI EG00229). All lungs perfused with PVS (PVS IRI) completed the perfusion protocol whereas all lungs perfused with PVS containing SU1498 (PVS IRI SU1498) reached the critical Pinsp (7.5mmHg) before the end of the experimental protocol (Figure 6A). In contrast, inhibition of VEGF-A binding (PVS IRI EG00229) had no effect on lung survival (Figure 6A). Pinsp when perfusion was stopped in the PVS IRI SU1498 group was significantly (P<0.001) higher than that in the PVS IRI or PVS IRI EG00229 groups (Figure 6D) and measurement of wet:dry weight ratios confirmed that oedema had occurred in the PVS IRI SU1498 group but not in the other two groups (Figure 6E). Capillary pressure was similar in all three groups (Figure 6F).

**Figure 6.**
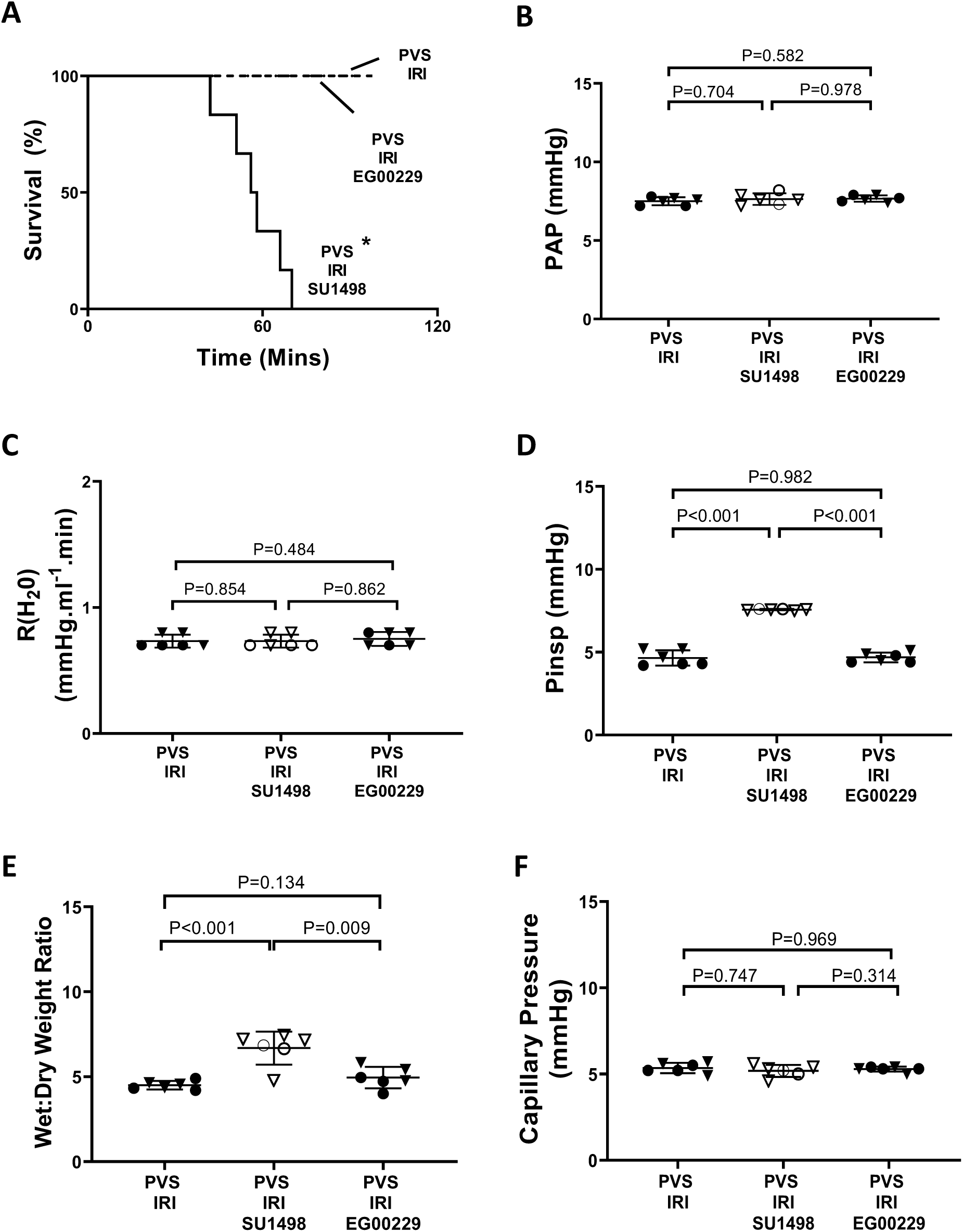
Protective effect of PVS is independent of VEGFA binding to VEGFR2-neuropilin 1 complex. A, Kaplan-Meier survival curves for injured lungs perfused with PVS (PVS IRI, n=6), injured lungs perfused with PVS with EG00229 added (PVS IRI EG00229, n=6) and injured lungs perfused with physiological viscosity solution with SU1498 added (PVS IRI SU1498, n=6). * indicates statistically significant difference from the two other groups (P<0.001, Log rank test with Holm-Sidak correction for multiple comparisons). B, mean (SD) pulmonary artery pressure (PAP) in groups at termination of perfusion (n=6 in each group). C, mean (SD) R(H_2_0) in groups at termination of perfusion (n=6 in each group). D, mean (SD) airway pressure (Pinsp) groups at termination of perfusion (n=6 in each group). E, mean (SD) wet to dry weight ratio in groups at termination of perfusion (n=6 in each group). F, mean (SD) capillary pressure in each group following reperfusion (n=6 in each group). Open symbols represent lungs in which perfusion was terminated due to the development of oedema (Pinsp>7.5mmHg) before 180 minutes. Filled symbols represent lungs in which oedema did not develop. Triangular symbols represent male lungs and circular symbols represent female lungs. Exact P values were computed using an un-paired t-test followed by Holm-Sidak correction for multiple comparisons. IRI, ischaemia reperfusion injury. LVS, low viscosity solution. Pinsp, peak inspiratory pressure. PAP, pulmonary arterial pressure. PVS, physiological viscosity solution. R(H_2_O), pulmonary vascular resistance corrected to perfusion with a solution whose relative viscosity equals one.

## Discussion

We report that in mouse lungs injured by ischaemia reperfusion injury perfusion with a solution that has reduced viscosity led to greater endothelial leak and oedema formation than perfusion at the same flow rate with a solution with a physiological viscosity i.e. a viscosity similar to that of blood *in vivo*. Since such perfusion conditions will alter shear stress on the endothelium, we investigated the role in this response of VEGFR2, an important endothelial mechanosensor. We found that the protective effects of the physiological viscosity solution were abolished by an inhibitor of autophosphorylation of the intracellular domain of VEGFR2 but were unaltered by an inhibitor of the binding of its ligand VEGFA to the extracellular domain.

IRI in the lung results in an acute lung injury characterised by endothelial damage and oedema formation (Zanotti, Casiraghi et al. 2009). In its early initiating phase the injury does not depend on the recruitment of leukocytes to the lung (Fiser, Tribble et al. 2001, Zhao, Fernandez et al. 2006). Our model of IRI recapitulated the key features of the early stages of IRI with progressively reducing dynamic compliance, increased wet-to-dry weight ratios and endothelial barrier leak, as previously reported (Fiser, Tribble et al. 2001, Zhao, Fernandez et al. 2006, Zanotti, Casiraghi et al. 2009). In addition, we showed that in the injured lungs oedema was caused by endothelial leak while capillary pressure remained unchanged. Thus, the oedema was not due to altered filtration pressure, a key feature of acute lung injury and adult respiratory distress syndrome in humans, which is by definition non-cardiogenic (Collins, Blank et al. 2013, Matthay, Zemans et al. 2019). It is interesting to note that this injury occurred even though ventilation continued with normoxic gas (21% O2) and thus the alveolar walls and capillaries had a continued supply of oxygen during the period of ischaemia (Chatterjee, Nieman et al. 2014). This is compatible with the loss of shear stress during the period of ischaemia being an important mechanism of injury. Using this model, we demonstrated that increasing shear stress, by increasing the viscosity of the perfusing solution to a level within the range observed in whole blood in vivo, at constant flow (Fahraeus RL 1931, Chien, Usami et al. 1966, Long, Smith et al. 2004), markedly reduced the rate of oedema formation by restoring endothelial barrier function toward normal i.e. the barrier function found in the uninjured lungs.

Previous *in vitro* experiments have shown that application of physiological levels of shear stress to endothelial cell monolayers from several different vascular beds causes cell alignment, increased inter-endothelial cell junctional protein and reduced permeability of the monolayers compared to cells cultured in the absence of shear stress (Seebach, Dieterich et al. 2000, Dekker, van Soest et al. 2002, Colgan, Ferguson et al. 2007, Siddharthan, Kim et al. 2007, Warboys, Eric Berson et al. 2010, Walsh, Murphy et al. 2011, Gross, Aggarwal et al. 2014, Kostyunina, Rowan et al. 2023). These reductions in permeability occur within hours of the onset of shear stress (Walsh, Murphy et al. 2011, Gross, Aggarwal et al. 2014, Rowan, Rochfort et al. 2018). Furthermore, the production of glycosaminoglycans, such as heparan sulphate, which are key structural components of the endothelial glycocalyx essential to normal barrier function, are also shear stress dependent (Torres Filho, Torres et al. 2013, Zeng and Tarbell 2014, Wang, Kostidis et al. 2020). Conversely, reduction of shear stress acting on monolayers previously conditioned by physiological shear stress caused an increase in permeability within hours of the reduction (Walsh, Murphy et al. 2011, Rowan, Rochfort et al. 2018). Thus, the rapid onset differences in permeability that we observed between lungs that were perfused with LVS and PVS are compatible with those previous in vitro findings. Our findings extend those previous observations by demonstrating for the first time that this beneficial effect of shear stress remains intact following IRI in the intact pulmonary circulation and can facilitate recovery from that injury.

While several mechanosensitive channels and receptors have been identified (Fels and Kusche-Vihrog 2020), one of the most important pathways implicated in endothelial specific mechano-transduction of shear stress is via a complex of proteins at the cell-cell junction consisting of the tyrosine kinase receptor vascular endothelial derived growth factor receptor 2 (VEGFR2), PECAM-1, and VE-cadherin (Jin, Ueba et al. 2003, Lee and Koh 2003, Coon, Baeyens et al. 2015, Mehta, Pang et al. 2020). Plexin D1 is a direct force sensor which forms the mechanosensitive complex with VEGFR2 and its co-receptor neuropilin 1 (NRP1) and causes activation of VEGFR2 in response to shear stress leading to autophosphorylation of the intracellular kinase insert domain and altered cellular function (Mehta, Pang et al. 2020). These responses include eNOS activation, cell alignment, flow-induced vasodilatation and a trophic action on endothelial cell in the normal systemic vasculature (Jin, Ueba et al. 2003, dela Paz, Walshe et al. 2012, Hein, Rosa et al. 2015). Inhibition of the tyrosine phosphorylation activity of the receptor with the selective small molecule inhibitor SU1498 blocked VEGFR2 autophosphorylation and the flow-induced vasodilation (Strawn, McMahon et al. 1996, Jin, Ueba et al. 2003, Tzima, Irani-Tehrani et al. 2005, Hein, Rosa et al. 2015). Our finding that SU1498 blocked the protective effect of the PVS following IRI provides evidence that the VEGFR2 receptor also has a central role in a shear stress induced enhancement of endothelial barrier function in the intact pulmonary circulation following injury. Furthermore, since in the presence of the inhibitor there was no longer any difference in endothelial leak and oedema formation between LVS and PVS solutions, our findings suggest that mechanotransduction via VEGFR2 mediated the entirety of the protective effect of PVS. This is compatible with the demonstration by Mehta and colleagues that ablation of the mechanoreceptor function of VEGFR2, by genetic deletion of plexinD1, completely abolished VEGFR2 mediated autophosphorylation and junctional complex assembly in response to shear stress in systemic endothelial cells (Mehta, Pang et al. 2020).

The canonical action of VEGF-A binding to VEGFR2 is to increase endothelial barrier permeability (Eliceiri, Paul et al. 1999, Carmeliet 2003) yet our data suggest an additional, important role for VEGFR2 in reducing endothelial permeability in the pulmonary vascular bed. It has previously been shown that shear stress-induced activation of VEGFR2 is independent of ligand binding (Jin, Ueba et al. 2003, Hein, Rosa et al. 2015) and that activation of VEGFR2 by shear stress initiates a downstream signalling cascade that is different from that produced by VEGF-A induced activation (Wang, Chang et al. 2007). Moreover, the induction of increased vascular permeability in response to VEGF-A can be blocked by mutation of a specific tyrosine phosphorylation site within VEGFR2 (Y949/Y951), while leaving the other trophic actions of VEGF-A intact (Li, Padhan et al. 2016, Smith, Ninchoji et al. 2020). Binding of VEGF-A to the coreceptor NRP1 together with the VEGFR2 homodimer is required for the VEGF-A induced increase in endothelial barrier permeability (Fantin et al. 2017). Thus, to exclude the possibility that VEGFA binding to VEGFR2 is involved in this effect of PVS, we used an inhibitor of VEGF-A binding to the VEGFR2-NRP1 co-receptor complex, EG00229 (Jarvis, Allerston et al. 2010, Fantin, Lampropoulou et al. 2017, Powell, Mota et al. 2018). We found that blockade of this pathway did not attenuate the action of PVS in reducing pulmonary oedema following IRI which, taken together with those previous reports, suggests that the protective effective effect of increased PVS was independent of VEGFA signalling.

It has previously been reported that VEGFR2 activates flow mediated vasodilatation in the ophthalmic circulation (Hein, Rosa et al. 2015). However, despite the potent effect of blockade of VEGFR2 autophosphorylation (by SU1498) on endothelial permeability that we observed, it had no effect on pulmonary vascular resistance. This suggests that other mechanoreceptors are responsible for flow mediated vasodilation in the pulmonary circulation. In support of this, evidence has been presented more recently that Piezo1 channels activate flow mediated vasodilatation in the pulmonary circulation (Lhomme, Gilbert et al. 2019, Porto Ribeiro, Barbeau et al. 2022).

Our findings potentially have important implications in clinical conditions which warrant further investigation in the future. VEGFA concentrations are increased in the plasma of patients with acute respiratory distress syndrome (ARDS) secondary to multiple different causes including COVID–19 (Thickett, Armstrong et al. 2001, Thickett, Armstrong and Millar 2002, Cross and Matthay 2011, Pang, Xu et al. 2021). In an *in vivo* animal model of ischaemia reperfusion-induced acute lung injury, blockade of activation of VEGFR2 by its ligand VEGFA using a monoclonal anti-VEGFA antibody attenuated oedema formation and lung injury (Lan, Peng et al. 2016) and, more recently, it has been reported that a monoclonal anti-VEGFA antibody reduced acute lung injury caused by COVID–19 and improved patient outcomes (Pang, Xu et al. 2021). Taken together with the results we report here, those previously published findings suggest a dual action of VEGFR2 in acute lung injury, a ligand-based activation promoting increased endothelial permeability and oedema formation and a mechanoreceptor mediated activation promoting reduced endothelial permeability. The net effect of VEGFR2 activation may then depend on the balance of these two activities in a particular circumstance. Since blockade of VEGFA binding to VEGFR2 leaves its mechanoreceptor function intact, as shown both by our data and previous reports (Jin, Ueba et al. 2003, Hein, Rosa et al. 2015), it may be the optimal strategy for targeting the VEGFA pathway in acute lung injury and ARDS. In contrast, blockers that target the tyrosine kinase activity of the intracellular domain of VEGFR2 should be avoided; this requires testing in further research. This may also be relevant to other clinical settings since ischaemia reperfusion injury (IRI) contributes to the pathophysiology of many common conditions, including myocardial infarction, pulmonary embolism, peripheral vascular insufficiency and hypovolemic shock (Grace 1994, Mongardon, Dumas et al. 2011) and IRI is also one of the principal determinants of organ failure following organ transplantation (Erasmus, van Raemdonck et al. 2016, Rampes and Ma 2019).

In summary, we have shown for the first time that the viscosity of the solution perfusing the pulmonary vascular bed following ischaemia-reperfusion injury is centrally important in determining the restoration and maintenance of normal endothelial barrier function. Furthermore, we found that this beneficial effect is abolished by a blocker of VEGFR2 autophosphorylation and downstream intracellular signalling. These findings have important implications for our understanding of tissue fluid balance and the mechanisms leading to non-cardiogenic oedema in acute lung injury and ARDS.

## Additional Information

### Data availability

Original data arising from this research are available directly from upon reasonable request.

### Competing interests

PMcL is a named co-inventor on a patent owned by University College Dublin, WO2016/162536, entitled “A lung perfusion solution, and use thereof for the ex vivo preservation of a mammalian lung,” publication date, October 23, 2016.

### Author contributions

All experimental work was undertaken in the Conway Institute of Biomolecular and Biomedical Research, University College Dublin.

DW, contributed to conception and design, data acquisition and analysis, data interpretation, drafting the manuscript and critically reviewing content. DK contributed to data acquisition and analysis, interpretation of data, critically reviewing and contributing to manuscript drafting. JB contributed to conception and design, interpretation of data, critically reviewing content and contributing to manuscript drafting. PMcL contributed to conception and design, data analysis and interpretation, drafting the manuscript and critically reviewing content. DW, DK, JB and PMcL have approved the final version of the manuscript, agree to be accountable for all aspects of the work in ensuring that questions related to the accuracy or integrity of any part of the work are appropriately investigated and resolved, agree that all persons designated as authors qualify for authorship, and all those who qualify for authorship are listed.

### Funding

British Journal of Anaesthesia/Royal College of Anaesthesia Project Grant WKR0-2019-0074, St Vincent’s Anaesthesia Foundation and Science Foundation Ireland 17/TIDA/4960.

## References

Aman, J., E. M. Weijers, G. P. van Nieuw Amerongen, A. B. Malik and V. W. van Hinsbergh (2016). “Using cultured endothelial cells to study endothelial barrier dysfunction: Challenges and opportunities.” Am J Physiol Lung Cell Mol Physiol 311(2): L453–466.

Annecke, T., J. Fischer, H. Hartmann, J. Tschoep, M. Rehm, P. Conzen, C. P. Sommerhoff and B. F. Becker (2011). “Shedding of the coronary endothelial glycocalyx: effects of hypoxia/reoxygenation vs ischaemia/reperfusion.” Br J Anaesth 107(5): 679–686.

Borek, I., A. Birnhuber, N. F. Voelkel, L. M. Marsh and G. Kwapiszewska (2023). “The vascular perspective on acute and chronic lung disease.” J Clin Invest 133(16).

Cadogan, E., N. Hopkins, S. Giles, J. G. Bannigan, J. Moynihan and P. McLoughlin (1999). “Enhanced expression of inducible nitric oxide synthase without vasodilator effect in chronically infected lungs.” American Journal of Physiology-Lung Cellular and Molecular Physiology 277(3): L616–L627.

Cahill, E., S. C. Rowan, M. Sands, M. Banahan, D. Ryan, K. Howell and P. McLoughlin (2012). “The pathophysiological basis of chronic hypoxic pulmonary hypertension in the mouse: vasoconstrictor and structural mechanisms contribute equally.” Exp Physiol 97(6): 796–806.

Carmeliet, P. (2003). “Angiogenesis in health and disease.” Nat Med 9(6): 653–660.

Chatpun, S. and P. Cabrales (2010). “Cardiac mechanoenergetic cost of elevated plasma viscosity after moderate hemodilution.” Biorheology 47(3-4): 225–237.

Chatterjee, S., G. F. Nieman, J. D. Christie and A. B. Fisher (2014). “Shear stress-related mechanosignaling with lung ischemia: lessons from basic research can inform lung transplantation.” Am J Physiol Lung Cell Mol Physiol 307(9): L668–680.

Chien, S., S. Usami, H. M. Taylor, J. L. Lundberg and M. I. Gregersen (1966). “Effects of hematocrit and plasma proteins on human blood rheology at low shear rates.” J Appl Physiol 21(1): 81–87.

Colgan, O. C., G. Ferguson, N. T. Collins, R. P. Murphy, G. Meade, P. A. Cahill and P. M. Cummins (2007). “Regulation of bovine brain microvascular endothelial tight junction assembly and barrier function by laminar shear stress.” American Journal of Physiology-Heart and Circulatory Physiology 292(6): H3190–H3197.

Collins, S. R., R. S. Blank, L. S. Deatherage and R. O. Dull (2013). “Special article: the endothelial glycocalyx: emerging concepts in pulmonary edema and acute lung injury.” Anesth Analg 117(3): 664–674.

Coon, B. G., N. Baeyens, J. Han, M. Budatha, T. D. Ross, J. S. Fang, S. Yun, J. L. Thomas and M. A. Schwartz (2015). “Intramembrane binding of VE-cadherin to VEGFR2 and VEGFR3 assembles the endothelial mechanosensory complex.” J Cell Biol 208(7): 975–986.

Cross, L. J. and M. A. Matthay (2011). “Biomarkers in acute lung injury: insights into the pathogenesis of acute lung injury.” Crit Care Clin 27(2): 355–377.

Davies, P. F. (1995). “Flow-mediated endothelial mechanotransduction.” Physiol Rev 75(3): 519–560.

Dekker, R. J., S. van Soest, R. D. Fontijn, S. Salamanca, P. G. de Groot, E. VanBavel, H. Pannekoek and A. J. Horrevoets (2002). “Prolonged fluid shear stress induces a distinct set of endothelial cell genes, most specifically lung Kruppel-like factor (KLF2).” Blood 100(5): 1689–1698.

Dekker, R. J., S. Van Soest, R. D. Fontijn, S. Salamanca, P. G. De Groot, E. VanBavel, H. Pannekoek and A. J. G. Horrevoets (2002). “Prolonged fluid shear stress induces a distinct set of endothelial cell genes, most specifically lung Krüppel-like factor (KLF2).” Blood 100: 1689–1698.

dela Paz, N. G., T. E. Walshe, L. L. Leach, M. Saint-Geniez and P. A. D’Amore (2012). “Role of shear-stress-induced VEGF expression in endothelial cell survival.” J Cell Sci 125(Pt 4): 831–843.

Eliceiri, B. P., R. Paul, P. L. Schwartzberg, J. D. Hood, J. Leng and D. A. Cheresh (1999). “Selective requirement for Src kinases during VEGF-induced angiogenesis and vascular permeability.” Mol Cell 4(6): 915–924.

Erasmus, M. E., D. van Raemdonck, M. Z. Akhtar, A. Neyrinck, D. G. de Antonio, A. Varela and J. Dark (2016). “DCD lung donation: donor criteria, procedural criteria, pulmonary graft function validation, and preservation.” Transpl Int 29(7): 790–797.

Fåhræus, R. and T. Lindqvist (1931). “THE VISCOSITY OF THE BLOOD IN NARROW CAPILLARY TUBES.” American Journal of Physiology-Legacy Content 96: 562-568.

Fahraeus RL, L. T. (1931). “The viscosity of blood in narrow capillary tubes.” Am J Physiol Heart Circ Physiol 96: 562–568

Fantin, A., A. Lampropoulou, V. Senatore, J. T. Brash, C. Prahst, C. A. Lange, S. E. Liyanage, C. Raimondi, J. W. Bainbridge, H. G. Augustin and C. Ruhrberg (2017). “VEGF165-induced vascular permeability requires NRP1 for ABL-mediated SRC family kinase activation.” J Exp Med 214(4): 1049–1064.

Fels, B. and K. Kusche-Vihrog (2020). “It takes more than two to tango: mechanosignaling of the endothelial surface.” Pflugers Arch 472(4): 419–433.

Fiser, S. M., C. G. Tribble, S. M. Long, A. K. Kaza, J. T. Cope, V. E. Laubach, J. A. Kern and I. L. Kron (2001). “Lung transplant reperfusion injury involves pulmonary macrophages and circulating leukocytes in a biphasic response.” J Thorac Cardiovasc Surg 121(6): 1069–1075.

Grace, P. A. (1994). “Ischaemia-reperfusion injury.” Br J Surg 81(5): 637–647.

Gross, C. M., S. Aggarwal, S. Kumar, J. Tian, A. Kasa, N. Bogatcheva, S. A. Datar, A. D. Verin, J. R. Fineman and S. M. Black (2014). “Sox18 preserves the pulmonary endothelial barrier under conditions of increased shear stress.” J Cell Physiol 229(11): 1802–1816.

Hausenloy, D. J., W. Chilian, F. Crea, S. M. Davidson, P. Ferdinandy, D. Garcia-Dorado, N. van Royen, R. Schulz and G. Heusch (2019). “The coronary circulation in acute myocardial ischaemia/reperfusion injury: a target for cardioprotection.” Cardiovasc Res 115(7): 1143–1155.

Hein, T. W., R. H. Rosa, Jr., Y. Ren, W. Xu and L. Kuo (2015). “VEGF Receptor-2-Linked PI3K/Calpain/SIRT1 Activation Mediates Retinal Arteriolar Dilations to VEGF and Shear Stress.” Invest Ophthalmol Vis Sci 56(9): 5381–5389.

Hoffman, J. I. (2011). “Pulmonary vascular resistance and viscosity: the forgotten factor.” Pediatr Cardiol 32(5): 557–561.

Jarvis, A., C. K. Allerston, H. Jia, B. Herzog, A. Garza-Garcia, N. Winfield, K. Ellard, R. Aqil, R. Lynch, C. Chapman, B. Hartzoulakis, J. Nally, M. Stewart, L. Cheng, M. Menon, M. Tickner, S. Djordjevic, P. C. Driscoll, I. Zachary and D. L. Selwood (2010). “Small Molecule Inhibitors of the Neuropilin-1 Vascular Endothelial Growth Factor A (VEGF-A) Interaction.” Journal of Medicinal Chemistry 53(5): 2215–2226.

Jin, Z. G., H. Ueba, T. Tanimoto, A. O. Lungu, M. D. Frame and B. C. Berk (2003). “Ligand-independent activation of vascular endothelial growth factor receptor 2 by fluid shear stress regulates activation of endothelial nitric oxide synthase.” Circ Res 93(4): 354–363.

Kloner, R. A., K. S. King and M. G. Harrington (2018). “No-reflow phenomenon in the heart and brain.” Am J Physiol Heart Circ Physiol 315(3): H550–H562.

Kostyunina, D. S., S. C. Rowan, N. V. Pakhomov, E. Dillon, K. D. Rochfort, P. M. Cummins, M. J. O’Rourke and P. McLoughlin (2023). “Shear Stress Markedly Alters the Proteomic Response to Hypoxia in Human Pulmonary Endothelial Cells.” Am J Respir Cell Mol Biol 68(5): 551–565.

Kwon, O. S., S. G. Noh, S. H. Park, R. H. I. Andtbacka, J. R. Hyngstrom and R. S. Richardson (2023). “Ageing and endothelium-mediated vascular dysfunction: the role of the NADPH oxidases.” J Physiol 601(3): 451–467.

Lan, C. C., C. K. Peng, S. E. Tang, S. Y. Wu, K. L. Huang and C. P. Wu (2016). “Anti-Vascular Endothelial Growth Factor Antibody Suppresses ERK and NF-kappaB Activation in Ischemia-Reperfusion Lung Injury.” PLoS One 11(8): e0159922.

Lee, H. J. and G. Y. Koh (2003). “Shear stress activates Tie2 receptor tyrosine kinase in human endothelial cells.” Biochem Biophys Res Commun 304(2): 399–404.

Levick, J. R. and C. C. Michel (2010). “Microvascular fluid exchange and the revised Starling principle.” Cardiovascular Research 87(2): 198–210.

Lhomme, A., G. Gilbert, T. Pele, J. Deweirdt, D. Henrion, I. Baudrimont, M. Campagnac, R. Marthan, C. Guibert, T. Ducret, J. P. Savineau and J. F. Quignard (2019). “Stretch-activated Piezo1 Channel in Endothelial Cells Relaxes Mouse Intrapulmonary Arteries.” Am J Respir Cell Mol Biol 60(6): 650–658.

Li, X., N. Padhan, E. O. Sjostrom, F. P. Roche, C. Testini, N. Honkura, M. Sainz-Jaspeado, E. Gordon, K. Bentley, A. Philippides, V. Tolmachev, E. Dejana, R. V. Stan, D. Vestweber, K. Ballmer-Hofer, C. Betsholtz, K. Pietras, L. Jansson and L. Claesson-Welsh (2016). “VEGFR2 pY949 signalling regulates adherens junction integrity and metastatic spread.” Nat Commun 7: 11017.

Long, D. S., M. L. Smith, A. R. Pries, K. Ley and E. R. Damiano (2004). “Microviscometry reveals reduced blood viscosity and altered shear rate and shear stress profiles in microvessels after hemodilution.” Proc Natl Acad Sci U S A 101(27): 10060–10065.

Ludbrook, J. (1998). “Multiple comparison procedures updated.” Clin Exp Pharmacol Physiol 25(12): 1032–1037.

Matthay, M. A., R. L. Zemans, G. A. Zimmerman, Y. M. Arabi, J. R. Beitler, A. Mercat, M. Herridge, A. G. Randolph and C. S. Calfee (2019). “Acute respiratory distress syndrome.” Nature Reviews Disease Primers 5(1): 18.

Mehta, D. and A. B. Malik (2006). “Signaling mechanisms regulating endothelial permeability.” Physiol Rev 86(1): 279–367.

Mehta, V., K.-L. Pang, D. Rozbesky, K. Nather, A. Keen, D. Lachowski, Y. Kong, D. Karia, M. Ameismeier, J. Huang, Y. Fang, A. del Rio Hernandez, J. S. Reader, E. Y. Jones and E. Tzima (2020). “The guidance receptor plexin D1 is a mechanosensor in endothelial cells.” Nature 578(7794): 290–295.

Mongardon, N., F. Dumas, S. Ricome, D. Grimaldi, T. Hissem, F. Pene and A. Cariou (2011). “Postcardiac arrest syndrome: from immediate resuscitation to long-term outcome.” Ann Intensive Care 1(1): 45.

Pak, O., A. Sydykov, D. Kosanovic, R. T. Schermuly, A. Dietrich, K. Schroder, R. P. Brandes, T. Gudermann, N. Sommer and N. Weissmann (2017). “Lung Ischaemia-Reperfusion Injury: The Role of Reactive Oxygen Species.” Adv Exp Med Biol 967: 195–225.

Pang, J., F. Xu, G. Aondio, Y. Li, A. Fumagalli, M. Lu, G. Valmadre, J. Wei, Y. Bian, M. Canesi, G. Damiani, Y. Zhang, D. Yu, J. Chen, X. Ji, W. Sui, B. Wang, S. Wu, A. Kovacs, M. Revera, H. Wang, X. Jing, Y. Zhang, Y. Chen and Y. Cao (2021). “Efficacy and tolerability of bevacizumab in patients with severe Covid-19.” Nat Commun 12(1): 814.

Pedersen, L., E. B. Nielsen, M. K. Christensen, M. Buchwald and M. Nybo (2014). “Measurement of plasma viscosity by free oscillation rheometry: imprecision, sample stability and establishment of a new reference range.” Ann Clin Biochem 51(Pt 4): 495–498.

Porto Ribeiro, T. S,. Barbeau, I. Baudrimont, P. Vacher, V. Freund-Michel, G. Cardouat, P. Berger, C. Guibert, T. Ducret and J. F. Quignard(2022). “Piezo1 Channel Activation Reverses Pulmonary Artery Vasoconstriction in an Early Rat Model of Pulmonary Hypertension: The Role of Ca(2+) Influx and Akt-eNOS Pathway.” Cells 11(15).

Powell, J., F. Mota, D. Steadman, C. Soudy, J. T. Miyauchi, S. Crosby, A. Jarvis, T. Reisinger, N. Winfield, G. Evans, A. Finniear, T. Yelland, Y. T. Chou, A. W. E. Chan, A. O’Leary, L. Cheng, D. Liu, C. Fotinou, C. Milagre, J. F. Martin, H. Jia, P. Frankel, S. Djordjevic, S. E. Tsirka, I. C. Zachary and D. L. Selwood (2018). “Small Molecule Neuropilin-1 Antagonists Combine Antiangiogenic and Antitumor Activity with Immune Modulation through Reduction of Transforming Growth Factor Beta (TGFbeta) Production in Regulatory T-Cells.” J Med Chem 61(9): 4135–4154.

Powell, J., F. Mota, D. Steadman, C. Soudy, J. T. Miyauchi, S. Crosby, A. Jarvis, T. Reisinger, N. Winfield, G. Evans, A. Finniear, T. Yelland, Y. T. Chou, A. W. E. Chan, A. O’Leary, L. Cheng, D. Liu, C. Fotinou, C. Milagre, J. F. Martin, H. Jia, P. Frankel, S. Djordjevic, S. E. Tsirka, I. C. Zachary and D. L. Selwood (2018). “Small Molecule Neuropilin-1 Antagonists Combine Antiangiogenic and Antitumor Activity with Immune Modulation through Reduction of Transforming Growth Factor Beta (TGFβ) Production in Regulatory T-Cells.” J Med Chem 61(9): 4135–4154.

Pries, A. R., T. W. Secomb and P. Gaehtgens (2000). “The endothelial surface layer.” Pflugers Arch 440(5): 653–666.

Rampes, S. and D. Ma (2019). “Hepatic ischemia-reperfusion injury in liver transplant setting: mechanisms and protective strategies.” J Biomed Res 33(4): 221–234.

Reitsma, S., D. W. Slaaf, H. Vink, M. A. M. J. Van Zandvoort and M. G. A. Oude Egbrink (2007). The endothelial glycocalyx: Composition, functions, and visualization. Pflugers Archiv European Journal of Physiology, Pflugers Arch. 454: 345–359.

Rowan, S. C., K. D. Rochfort, L. Piouceau, P. M. Cummins, M. O’Rourke and P. McLoughlin (2018). “Pulmonary endothelial permeability and tissue fluid balance depend on the viscosity of the perfusion solution.” American Journal of Physiology-Lung Cellular and Molecular Physiology 315: L476–L484.

Schmidt, E. P., Y. Yang, W. J. Janssen, A. Gandjeva, M. J. Perez, L. Barthel, R. L. Zemans, J. C. Bowman, D. E. Koyanagi, Z. X. Yunt, L. P. Smith, S. S. Cheng, K. H. Overdier, K. R. Thompson, M. W. Geraci, I. S. Douglas, D. B. Pearse and R. M. Tuder (2012). “The pulmonary endothelial glycocalyx regulates neutrophil adhesion and lung injury during experimental sepsis.” Nat Med 18(8): 1217–1223.

Seebach, J., P. Dieterich, F. Luo, H. Schillers, D. Vestweber, H. Oberleithner, H.-J. Galla and H.-J. Schnittler (2000). “Endothelial barrier function under laminar fluid shear stress.” Laboratory investigation 80(12): 1819–1831.

Siddharthan, V., Y. V. Kim, S. Liu and K. S. Kim (2007). “Human astrocytes/astrocyte-conditioned medium and shear stress enhance the barrier properties of human brain microvascular endothelial cells.” Brain research 1147: 39–50.

Siddharthan, V., Y. V. Kim, S. Liu and K. S. Kim (2007). “Human astrocytes/astrocyte-conditioned medium and shear stress enhance the barrier properties of human brain microvascular endothelial cells.” Brain Res 1147: 39–50.

Smith, R. O., T. Ninchoji, E. Gordon, H. Andre, E. Dejana, D. Vestweber, A. Kvanta and L. Claesson-Welsh (2020). “Vascular permeability in retinopathy is regulated by VEGFR2 Y949 signaling to VE-cadherin.” Elife 9.

Snow, H. M., F. Markos, D. O’Regan and K. Pollock (2001). “Characteristics of arterial wall shear stress which cause endothelium-dependent vasodilatation in the anaesthetized dog.” J Physiol 531(Pt 3): 843–848.

Strawn, L. M., G. McMahon, H. App, R. Schreck, W. R. Kuchler, M. P. Longhi, T. H. Hui, C. Tang, A. Levitzki, A. Gazit, I. Chen, G. Keri, L. Orfi, W. Risau, I. Flamme, A. Ullrich, K. P. Hirth and L. K. Shawver (1996). “Flk-1 as a target for tumor growth inhibition.” Cancer Res 56(15): 3540–3545.

Thickett, D. R., L. Armstrong, S. J. Christie and A. B. Millar (2001). “Vascular endothelial growth factor may contribute to increased vascular permeability in acute respiratory distress syndrome.” Am J Respir Crit Care Med 164(9): 1601–1605.

Thickett, D. R., L. Armstrong and A. B. Millar (2002). “A role for vascular endothelial growth factor in acute and resolving lung injury.” Am J Respir Crit Care Med 166(10): 1332–1337.

Torres Filho, I. L,. N. Torres, J. L. Sondeen, I. A. Polykratis and M. A. Dubick (2013). “In vivo evaluation of venular glycocalyx during hemorrhagic shock in rats using intravital microscopy.” Microvasc Res 85: 128–133.

Townsley, M. I., R. J. Korthuis, B. Rippe, J. C. Parker and A. E. Taylor (1986). “Validation of double vascular occlusion method for Pc,i in lung and skeletal muscle.” J Appl Physiol (1985) 61(1): 127–132.

Tzima, E., M. Irani-Tehrani, W. B. Kiosses, E. Dejana, D. A. Schultz, B. Engelhardt, G. Cao, H. DeLisser and M. A. Schwartz (2005). “A mechanosensory complex that mediates the endothelial cell response to fluid shear stress.” Nature 437(7057): 426–431.

van Haaren, P. M., E. VanBavel, H. Vink and J. A. Spaan (2003). “Localization of the permeability barrier to solutes in isolated arteries by confocal microscopy.” Am J Physiol Heart Circ Physiol 285(6): H2848–2856.

Vaya, A., M. Simo, M. Santaolaria, P. Carrasco and D. Corella (2007). “Plasma viscosity and related cardiovascular risk factors in a Spanish Mediterranean population.” Thromb Res 120(4): 489–495.

Walsh, T. G., R. P. Murphy, P. Fitzpatrick, K. D. Rochfort, A. F. Guinan, A. Murphy and P. M. Cummins (2011). “Stabilization of brain microvascular endothelial barrier function by shear stress involves VE-cadherin signaling leading to modulation of pTyr-occludin levels.” Journal of cellular physiology 226(11): 3053–3063.

Wang, G., S. Kostidis, G. L. Tiemeier, W. M. P. J. Sol, M. R. de Vries, M. Giera, P. Carmeliet, B. M. van den Berg and T. J. Rabelink (2020). “Shear Stress Regulation of Endothelial Glycocalyx Structure Is Determined by Glucobiosynthesis.” Arteriosclerosis, thrombosis, and vascular biology 40: 350–364.

Wang, Y., J. Chang, K.-D. Chen, S. Li, J. Y.-S. Li, C. Wu and S. Chien (2007). “Selective adapter recruitment and differential signaling networks by VEGF vs. shear stress.” Proceedings of the National Academy of Sciences 104(21): 8875–8879.

Warboys, C. M., R. Eric Berson, G. E. Mann, J. D. Pearson and P. D. Weinberg (2010). “Acute and chronic exposure to shear stress have opposite effects on endothelial permeability to macromolecules.” Am J Physiol Heart Circ Physiol 298(6): H1850–1856.

Yin, J., L. Michalick, C. Tang, A. Tabuchi, N. Goldenberg, Q. Dan, K. Awwad, L. Wang, L. Erfinanda, G. Nouailles, M. Witzenrath, A. Vogelzang, L. Lv, W. L. Lee, H. Zhang, O. Rotstein, A. Kapus, K. Szaszi, I. Fleming, W. B. Liedtke, H. Kuppe and W. M. Kuebler (2016). “Role of Transient Receptor Potential Vanilloid 4 in Neutrophil Activation and Acute Lung Injury.” Am J Respir Cell Mol Biol 54(3): 370–383.

Zanotti, G., M. Casiraghi, J. B. Abano, J. R. Tatreau, M. Sevala, H. Berlin, S. Smyth, W. K. Funkhouser, K. Burridge, S. H. Randell and T. M. Egan (2009). “Novel critical role of Toll-like receptor 4 in lung ischemia-reperfusion injury and edema.” Am J Physiol Lung Cell Mol Physiol 297(1): L52–63.

Zeng, Y. and J. M. Tarbell (2014). “The adaptive remodeling of endothelial glycocalyx in response to fluid shear stress.” PLoS One 9(1): e86249.

Zhao, M., L. G. Fernandez, A. Doctor, A. K. Sharma, A. Zarbock, C. G. Tribble, I. L. Kron and V. E. Laubach (2006). “Alveolar macrophage activation is a key initiation signal for acute lung ischemia-reperfusion injury.” Am J Physiol Lung Cell Mol Physiol 291(5): L1018–1026.

